# An in-depth look at DNA crystals through the prism of molecular dynamics simulations

**DOI:** 10.1101/413336

**Authors:** Antonija Kuzmanic, Pablo D. Dans, Modesto Orozco

## Abstract

X-ray crystallography has been traditionally considered as the primary tool for the determination of biomolecular structures and its derived models are taken as the gold standard in structural biology. However, contacts formed through the crystal lattice are known to affect the structures, especially in the case of small and flexible molecules, like DNA oligos, introducing drastic changes in the structure with respect to the solution phase. Furthermore, it is still unknown why molecules crystallize in certain symmetry groups and how the associated lattice impacts their structure. The role of crystallization additives and whether they are just innocuous and unspecific catalyzers of the crystallization process also remains unclear. On account of a massive computational effort and the use of the latest generation force field, we were able to describe with unprecedented level of detail the nature of intermolecular forces that participate in the stabilization of B-DNA crystals in various symmetry groups and in different solvent environments. We showed that the stability of the crystal lattice and the type of crystallization additives are tightly coupled, and certain symmetry groups are only stable in the presence of a specific crystallization additive (i.e., spermine). Additives and crystal contacts induce small but non-negligible changes in the physical properties of DNA.

## INTRODUCTION

Ever since Linus Pauling laid the foundations of structural biology, X-ray crystallography has been the cornerstone method for solving biomolecular 3D structures. Hundreds of thousands of models derived from X-ray data and deposited in the Protein Data Bank (PDB) are the best witnesses of the power of this technique, which has also become the gold standard for validating other structural methods. However, just as any other method, X-ray crystallography has its own shortcomings and obtaining diffractable crystals is not always an easy task.^1–3^ Through robotic equipment, numerous variants of crystallization buffers are tested until a suitable one is found.^4^ However, why a given buffer promotes crystallization, what its influence is on the symmetry of the unit cell, as well as the overall structure and physical properties of the biomolecule, is typically unknown.

Thousands of X-ray derived DNA structures have been crucial in understanding the fine details of isolated and protein-bound DNAs.^5–7^ Unfortunately, artifacts in DNA crystals can lead to conformations which are otherwise undetected in solution (like the A-form of DNA). Even in ideal cases, such as the Drew-Dickerson dodecamer (DDD^8,9^), where crystal structures resemble the solution ones reasonably well, it is unclear how crystallization conditions modulate the symmetry space in which the molecule crystalizes, or the fine details of the structure and the physical properties of the DNA.^1,2^ Knowing the molecular interactions that stabilize the crystals would help to understand the actual properties of DNA in such systems and the mechanisms through which highly charged molecules are packed in severely crowded conditions, e.g. the cellular nucleus.

Here we use a new generation of supercomputers and a recently developed DNA force field^10–12^ to analyze the nature of DDD in three known crystal lattices and to explore the effects of crystallization additives on the stability of the crystals and the properties of the DNA.^13–15^ Through extensive unbiased MD simulations on the multi-microsecond timescale, we demonstrate how the stability of DNA crystals depends on subtle interactions between the packed DNA molecules and the components of the crystallization buffer. We obtained, for the first time, stable atomistic simulations of DNA crystals in biologically relevant timescales,^16–21^ in various symmetry groups (in which DDD has been crystallized), and in different solvent environments, which allowed us to understand with unprecedented level of detail the nature of intermolecular interactions that stabilize the crystals. Through the analysis of numerous simulations, we characterized the physical properties of DNA in a highly crowded environment resembling chromatin, and how the presence of additives affects them.

## METHODS

### Systems setup

X-ray structures of the Drew-Dickerson dodecamer used as the starting conformations in the unbiased MD simulations are those deposited under PDB codes 1EHV,^14^ 1FQ2,^13^ and 463D.^15^ Both 1EHV and 1FQ2 entries contain the full DNA duplex, while the 5’ cytosines are missing in 463D and were modelled in with PyMOL v1.8.2^22^ by using 1EHV as the template. The dodecahedral simulation boxes for solution systems (labelled sol) were created by solvating a single DNA duplex with TIP3P water molecules,^23,24^ leaving a 10-Å solvent layer around the duplex (∼20,000 atoms). K^+^, Na^+^, and Cl^-^ ions (Smith and Dang parameters)^25,26^ were added to reach neutrality and the final concentration of 400 mM (with 4:3 potassium to sodium ratio). In the case of 1FQ2 1:6 sol system, 6 spermine molecules were randomly added prior to solvation with TIP3P water and neutrality was reached with Cl^-^ ions. For the crystal systems, the simulation boxes were built following the dimensions of the unit cells reported in the PDB using PyMOL v1.8. The Na/KCl and MgCl2 systems were completed by first adding TIP3P water and then the ions to reach neutrality and the final concentrations listed in Table 1 (using Smith and Dang^25,26^ parameters for Na/KCl systems and Allnér-Nilsson-Villa^27^ parameters for MgCl_2_). The SpmCl_4_ and MgCl_2_·6H_2_O crystal systems were completed by first randomly adding spermine or hexacoordinated Mg^2+^ ions (using Yoo and Aksimentiev^28,29^ parameters, quantities in Table 1), then TIP3P waters followed by monovalent counter ions. System sizes ranged from 44,000-60,000 atoms for the smaller crystals, while the 1EHV systems with 81 duplexes had ∼170,000 atoms.

**Table 1.** Studied systems and simulated conditions.

| System name | PDB | Environment | Space group | Nº of unit cells | Nº of DNA duplexes | Salt type | [Salt] (mM) | dsDNA to Spm ratio | Temp. (K) | Equilibration time (ps) | Simulation time (μs) |
| --- | --- | --- | --- | --- | --- | --- | --- | --- | --- | --- | --- |
| 1EHV sol | 1EHV | solution | --- | --- | 1 | Na/KCl | 400 | --- | 284 | 600 | 1 |
| 15 mM | 1EHV | crystal | P3<sub>2</sub>12 | 3x3x3 | 81 | Na/KCl | 15 | --- | 284 | 600 | 1 |
| 200 mM | 1EHV | crystal | P3<sub>2</sub>12 | 3x3x3 | 81 | Na/KCl | 200 | --- | 284 | 600 | 1 |
| 400 mM | 1EHV | crystal | P3<sub>2</sub>12 | 3x3x3 | 81 | Na/KCl | 400 | --- | 284 | 600 | 4 |
| 600 mM | 1EHV | crystal | P3<sub>2</sub>12 | 3x3x3 | 81 | Na/KCl | 600 | --- | 284 | 600 | 1 |
| 800 mM | 1EHV | crystal | P3<sub>2</sub>12 | 3x3x3 | 81 | Na/KCl | 800 | --- | 284 | 600 | 1 |
| 1EHV Na/KCl | 1EHV | crystal | P3<sub>2</sub>12 | 3x3x1 | 27 | Na/KCl | 400 | --- | 284 | 600 | 4 |
| 1EHV Mg | 1EHV | crystal | P3<sub>2</sub>12 | 3x3x1 | 27 | MgCl<sub>2</sub> | 400 | --- | 284 | 600 | 4 |
| 1EHV Mg hex | 1EHV | crystal | P3<sub>2</sub>12 | 3x3x1 | 27 | MgCl<sub>2</sub>·6H<sub>2</sub>O | 400 | --- | 284 | 600 | 4 |
| 1EHV 1:6 | 1EHV | crystal | P3<sub>2</sub>12 | 3x3x1 | 27 | SpmCl<sub>4</sub> | --- | 1:6 | 284 | 6000 | 1 |
| 1EHV 1:9 | 1EHV | crystal | P3<sub>2</sub>12 | 3x3x1 | 27 | SpmCl<sub>4</sub> | --- | 1:9 | 284 | 6000 | 1 |
| 1EHV 1:12 | 1EHV | crystal | P3<sub>2</sub>12 | 3x3x1 | 27 | SpmCl<sub>4</sub> | --- | 1:12 | 284 | 6000 | 1 |
| 1FQ2 sol | 1FQ2 | solution | --- | --- | 1 | Na/KCl | 400 | --- | 295 | 600 | 1 |
| 1FQ2 1:6 sol | 1FQ2 | solution | --- | --- | 1 | SpmCl<sub>4</sub> | --- | 1:6 | 295 | 600 | 1 |
| 1FQ2 Na/KCl | 1FQ2 | crystal | P2<sub>1</sub>2<sub>1</sub>2<sub>1</sub> | 3x2x1 | 24 | Na/KCl | 400 | --- | 295 | 600 | 4 |
| 1FQ2 Na/KCl equ | 1FQ2 | crystal | P2<sub>1</sub>2<sub>1</sub>2<sub>1</sub> | 3x2x1 | 24 | Na/KCl | 400 | --- | 295 | 6000 | 1 |
| 1FQ2 1:3 | 1FQ2 | crystal | P2<sub>1</sub>2<sub>1</sub>2<sub>1</sub> | 3x2x1 | 24 | SpmCl<sub>4</sub> | --- | 1:3 | 295 | 6000 | 1 |
| 1FQ2 1:6 | 1FQ2 | crystal | P2<sub>1</sub>2<sub>1</sub>2<sub>1</sub> | 3x2x1 | 24 | SpmCl<sub>4</sub> | --- | 1:6 | 295 | 6000 | 1 |
| 1FQ2 1:9 | 1FQ2 | crystal | P2<sub>1</sub>2<sub>1</sub>2<sub>1</sub> | 3x2x1 | 24 | SpmCl<sub>4</sub> | --- | 1:9 | 295 | 6000 | 1 |
| 463D sol | 463D | solution | --- | --- | 1 | Na/KCl | 400 | --- | 293 | 600 | 1 |
| 463D | 463D | crystal | H3 | 2x2x1 | 36 | Na/KCl | 400 | --- | 293 | 6000 | 4 |

### Relaxation and equilibration of the systems

Energy of the simulated systems (listed in Table 1) was initially minimized through steepest-descent energy minimization. The initial velocities for the atoms were taken from Maxwell distribution at 100 K and the systems were subsequently heated to the final temperature (corresponding to the temperatures at which the crystals were grown; Table 1) in five steps of ∼50 K simulated for either 100 or 1000 ps at constant volume and temperature using velocity rescale thermostat.^30^ In parallel, atomic position restraints for the DNA molecules were uniformly relaxed. To achieve internal pressure close to 1 bar in crystal simulations, additional TIP3P water molecules were randomly placed within each system, which was first minimized and equilibrated, and then 50-ns production simulations were run, in a trial and error process until a suitable pressure was obtained (see Supp. Methods for further details).

### Production trajectories

Once simulation conditions matched those expected in the crystal or in solution, production runs (1-4 μs long) were generated using GROMACS 5.0.4 biomolecular simulation package^31^ with a 2-fs integration step and coordinate output of 10 ps. Constant volume conditions were employed for the crystal simulations, while the constant pressure (1 bar) was used in solution simulations. All simulations were done at constant temperature (284-295 K depending on the reported crystallization conditions). Parrinello-Rahman barostat^32^ and velocity rescale thermostat were used.^30^ Periodic boundary conditions and particle mesh Ewald^33^ were used in conjunction with state-of-the-art simulation conditions. DNA was described using the recently developed parmbsc1 force field.^10^ See Methods for additional details of the simulations.

### Analysis

Center of mass (COM) displacements were calculated after the optimal roto-translational fitting of the whole system to the ideal crystal. RMSD values were calculated for each duplex after the optimal roto-translational fitting to the original X-ray structures (see Suppl. Methods for details). Diffusion coefficients were calculated from a linear fit of mean square displacements over time using Einstein’s equation. The effective temperature of water molecules in crystals was estimated from the corresponding diffusion coefficients which were first scaled by 1/2.56 (to account for the overestimation of the water diffusion by the TIP3P water model) as described in our previous work,^34^ while effective DNA temperature was inferred at the base pair step level from the analysis of helical covariance matrices, which provided the stiffness matrices associated to harmonic deformations in the helical space.^35^ Essential dynamics was determined by the diagonalization of the Cartesian covariance matrices. Similarity in essential deformation modes were computed using Hess^36^ and Pérez *et al*. metrics,^35^ without considering the capping base pairs. Additional details on the analysis and the associated packages are shown in Suppl. Methods.

### Data availability

All trajectories reported here are deposited in the MuG-BigNASim^37^ database and can be freely downloaded. http://www.multiscalegenomics.eu/MuGVRE/modules/BigNASimMuG/

## RESULTS AND DISCUSSION

### Are the simulations of DNA crystals stable in the μs regime?

The Drew-Dickerson dodecamer is a prototypical B-DNA molecule, whose structure has been determined by X-ray crystallography in various space groups and under diverse conditions,^13–15^ as well as through high-quality NMR data in solution. We chose to perform our simulations on structures crystallized in 3 different space groups: P3212, P212121, and H3 (PDB codes: 1EHV, 1FQ2, and 463D, respectively). We followed the multistep protocol developed by the Case group,^17^ which includes a careful stabilization of the internal pressure by calibrating the number of water molecules in the system (see the Supp. Methods section for details and Supp. Table S1). Once the internal pressure was adjusted, we ran multi-microsecond long simulations of each crystal, which contained respectively 27, 24, and 36 copies of DDD (Table 1). To our surprise, two of our systems showed a systematic degradation of the crystal lattice, as shown by the center-of-mass (COM) displacements of each duplex with respect to the ideal positions in the crystal (Figure 1). The distortion of the lattice was particularly pronounced along the z-axis and got worse with simulation time for 1EHV and 1FQ2, while the 463D system was completely stable thorough the trajectory. Not only was the lattice intact in 463D, but the MD conformations sampled in the helical space on the base pair step (bps) level were in complete agreement with the average X-ray values (Figure 2), thus establishing that our force field is fully capable of representing local and global features of crystal structures.

**Figure 1.**
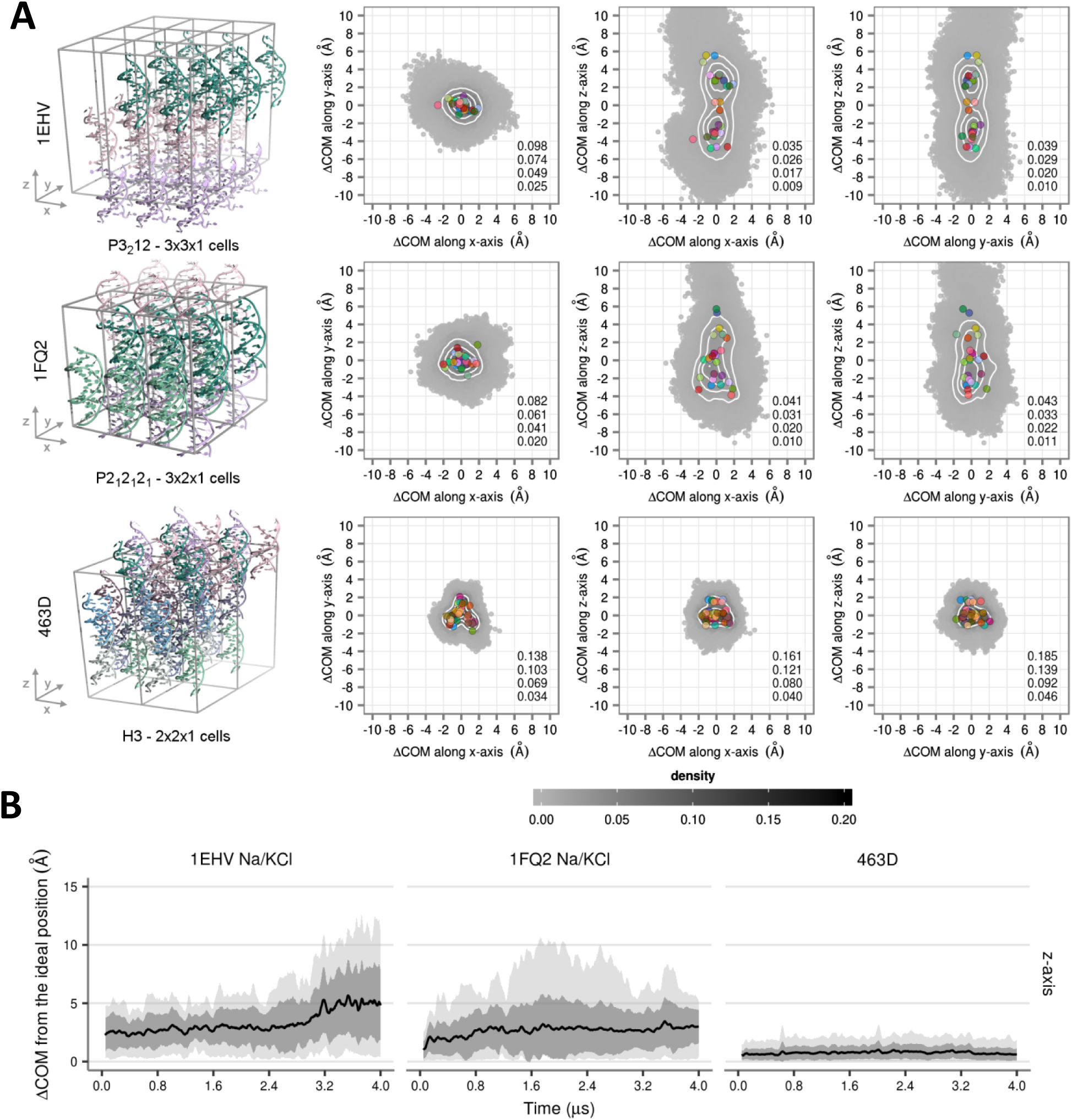
A) COM displacements from the ideal crystallographic positions shown for each duplex in the simulated supercell systems (1EHV, 1FQ2, 463D) along principal axes. Points are colored in greyscale according to the smooth 2D densities, estimated by fitting the observed distributions to a bivariate normal kernel (evaluated on a square grid of 90 × 90 bins). Four isodensity curves are shown in white and are quantified on the bottom right side of each plot. The average positions of the duplexes are given as colored circles. B) COM displacements in time along the z-axis shown for all three systems. Mean values are shown as black lines, standard deviations are shown as dark grey ribbons, while the extreme values are shown with light grey ribbons (lower panels).

**Figure 2.**
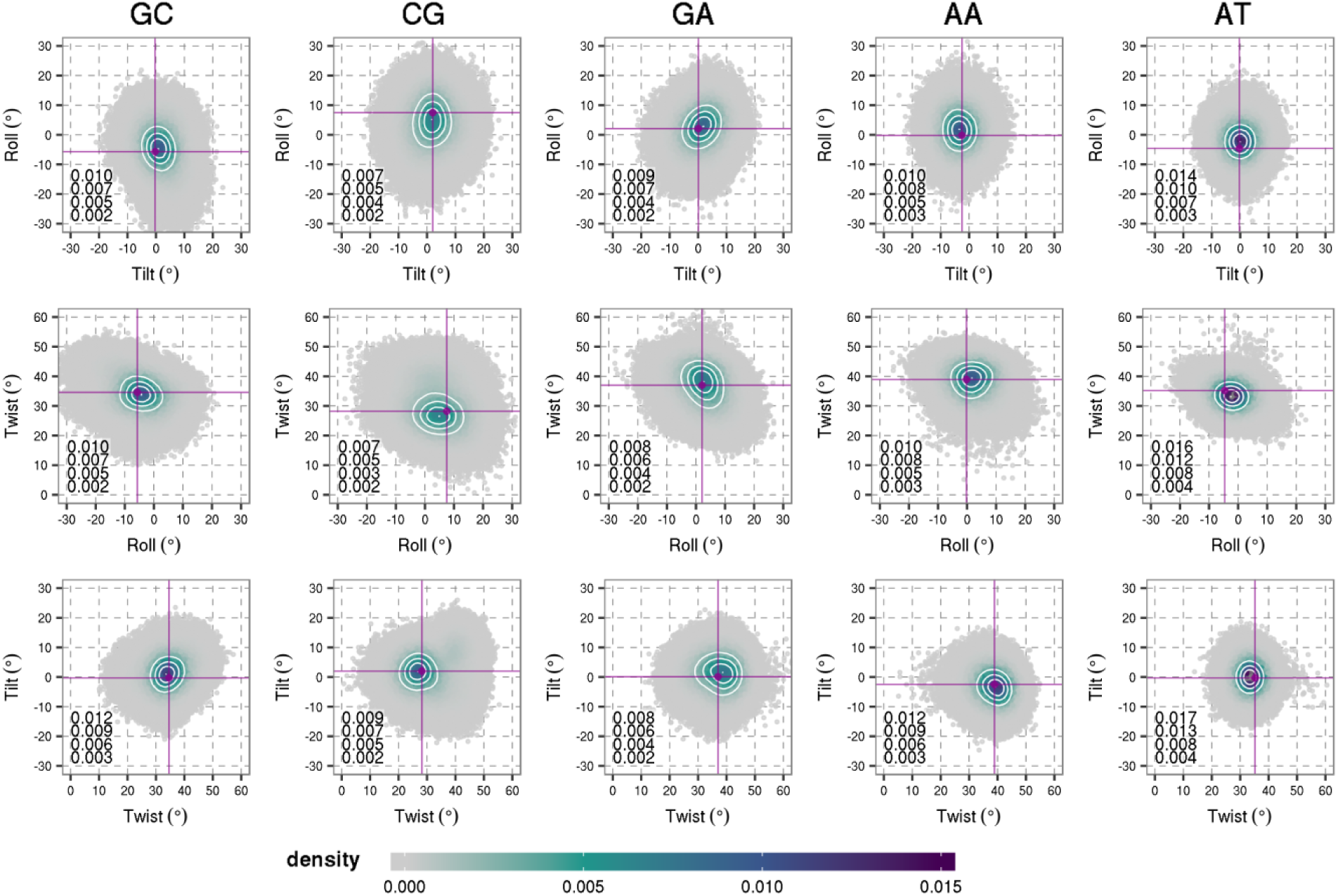
Rotational helical parameters (roll, twist, tilt) of all the duplexes in the 463D system (given for distinct base pair steps in DDD after removing the ends). The values calculated from the X-ray structure are shown with purple lines. Points are colored according to the smooth 2D densities, estimated by fitting the observed distributions to a bivariate normal kernel (evaluated on a square grid of 90 × 90 bins). Four isodensity curves are shown in white and are quantified on the bottom right side of each plot.

We thus centered our efforts on stabilizing the other two crystals starting first with the smallest system: 1EHV. We simulated a larger crystal (81 instead of 27 DDD molecules) with 5 different Na/KCl concentrations (Table 1), ranging from 15 mM to 800 mM (i.e., up to 5 times the assumed physiological salt concentration of the cellular nucleus), without being able to stabilize it (Supp. Figure S1). Considering that the increase in crystal’s size had little effect on its stability (Supp. Figure S2), we decided to proceed with the smaller system (with 27 DDD copies). The addition of 400 mM MgCl2 produced some improvement in the lattice integrity, but its degradation was still evident during the 4-μs-long simulations (Supp. Figure S3, and Supp. Methods for more details). We thus checked if the global degradation of the crystals was due to internal distortions of the double helices by analyzing the 2D distributions of the bps helical parameters from MD simulations. We obtained a good agreement between the values calculated from MD simulations and the X-ray structures (Supp. Figure S4), which discards the notion that the lattice distortions were caused by the deterioration of duplex geometries. Even the flexible ends (residue pairs 1-24 and 12-13), which exhibit fast but moderate fraying in solution,^11^ had in most cases an average RMSD with respect to the X-ray structures under 4 Å, oscillating between 3D conformations compatible with the experimental electron density (Supp. Figure S5). We obtained similar results when trying to stabilize, without success, the P2_1_2_1_2_1_ 1FQ2 crystal (data not shown), which allowed us to discard the possibility of a force field artifact that is space-group specific.

### How important are the buffer components in the stabilization of DNA crystals?

As we had discarded any obvious explanation for the degradation of the crystal lattice, we focused our attention on the chemical additives used to obtain the DDD crystals in the studied space groups. We noticed that 463D system was crystallized without using spermine (SPM),^15^ while both 1EHV^14^ and 1FQ2^13^ crystals were formed in the presence of this polyamine (+4 charge at pH 7). Spermine is normally found in all eukaryotic cells and has been used as an “inert molecular glue” for obtaining thousands of DNA crystals in all the major forms (A, B, Z, etc.).^19,20,38^ However, the molecular basis of its mechanism of action is mostly unknown, and in fact, spermine electron densities are often absent from the crystal. To check whether the lack of spermine in our simulations was the reason of the degradation of P3212 and P212121 simulations, we repeated the simulations for both crystals at three different spermine concentrations ranging from 1:3 (3 spermine molecules per duplex) up to 1:12 (Table 1). At last, the lattice integrity for both space groups was now preserved in a consistent concentration dependent manner (Figure 3). At the medium spermine concentration tested, 1:9 for 1EHV and 1:6 for 1FQ2, the observed stability is already comparable to the one obtained for 463D with no perceptible drift of the crystal lattice along the y/z-axes (Supp. Figures S6 and S7). Clearly the “inert molecular glue” has a major role in preserving the integrity of some of the crystal lattices.

**Figure 3.**
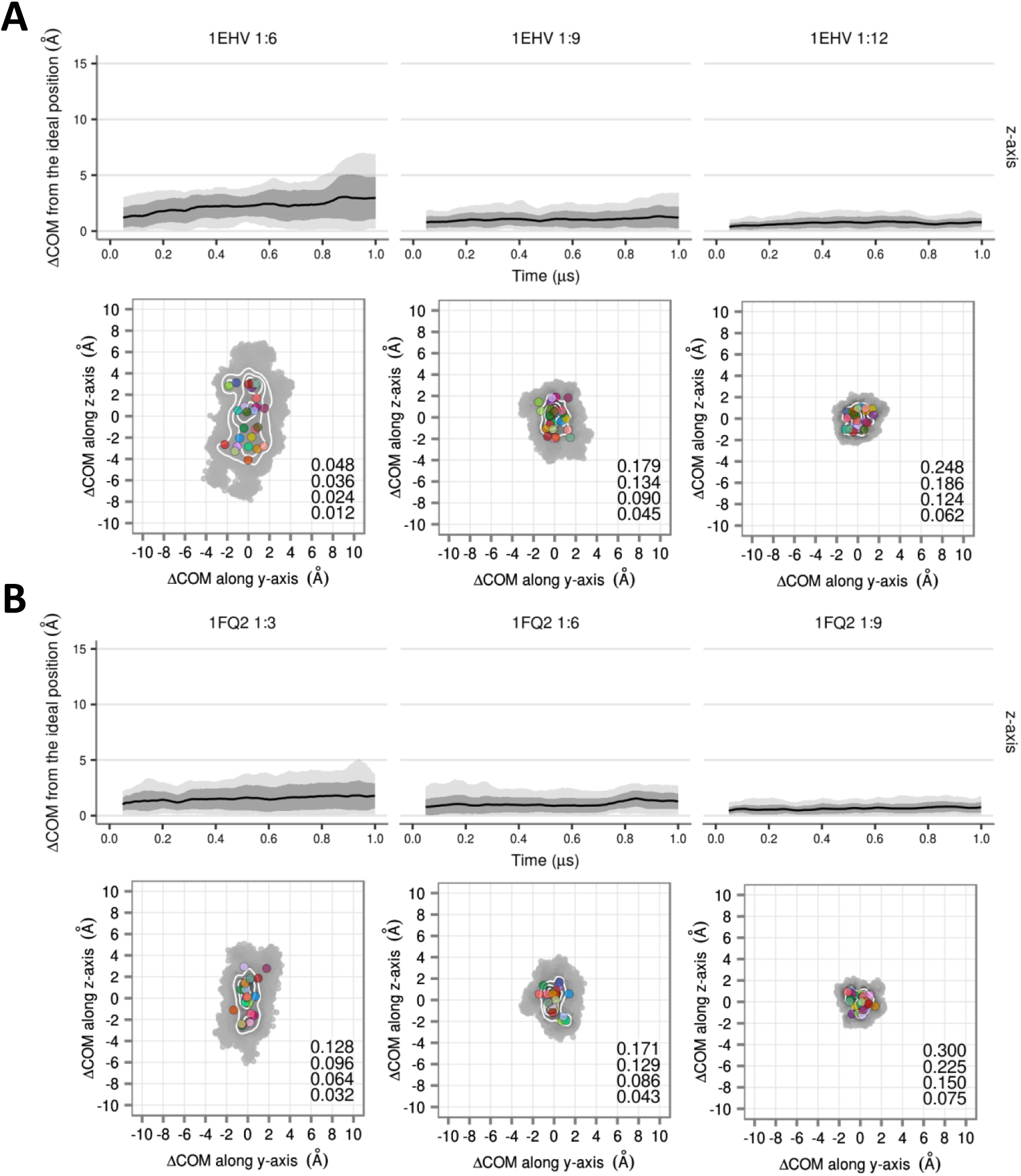
COM displacements from the ideal crystallographic positions for different spermine concentrations in A) 1EHV and B) 1FQ2 systems. The upper panels show the COM displacements in time along the z-axis where mean values are shown as black lines, standard deviations are shown as dark grey ribbons, while the extreme values are shown with light grey ribbons. The lower panels show COM displacements shown for each duplex along y and z axes with the average positions of the duplexes given as colored circles. Points are colored in greyscale according to the smooth 2D densities, estimated by fitting the observed distributions to a bivariate normal kernel (evaluated on a square grid of 90 × 90 bins). Four isodensity curves are shown in white and are quantified on the bottom right side of each plot.

Similarly to 463D, once properly stabilized, we obtained an impressive agreement between the X-ray structures (1EHV/1FQ2) and the MD crystals. This is visible when comparing the rotational helical space (Figure 4), or the groove dimensions, a property which has always been difficult to reproduce accurately by modern force fields (Supp. Figure S8). The dynamics of the end residues also seems to be stable according to their RMSD values, which are typically below 2 Å with respect to the crystal position (Supp. Figures S9 and S10). In the end, to illustrate the observed structural agreement, we also compared the average structures obtained from crystal MD simulations with the deposited X-ray structure for the three systems stabilized in the μs regime is shown in Supp. Figure S11.

**Figure 4.**
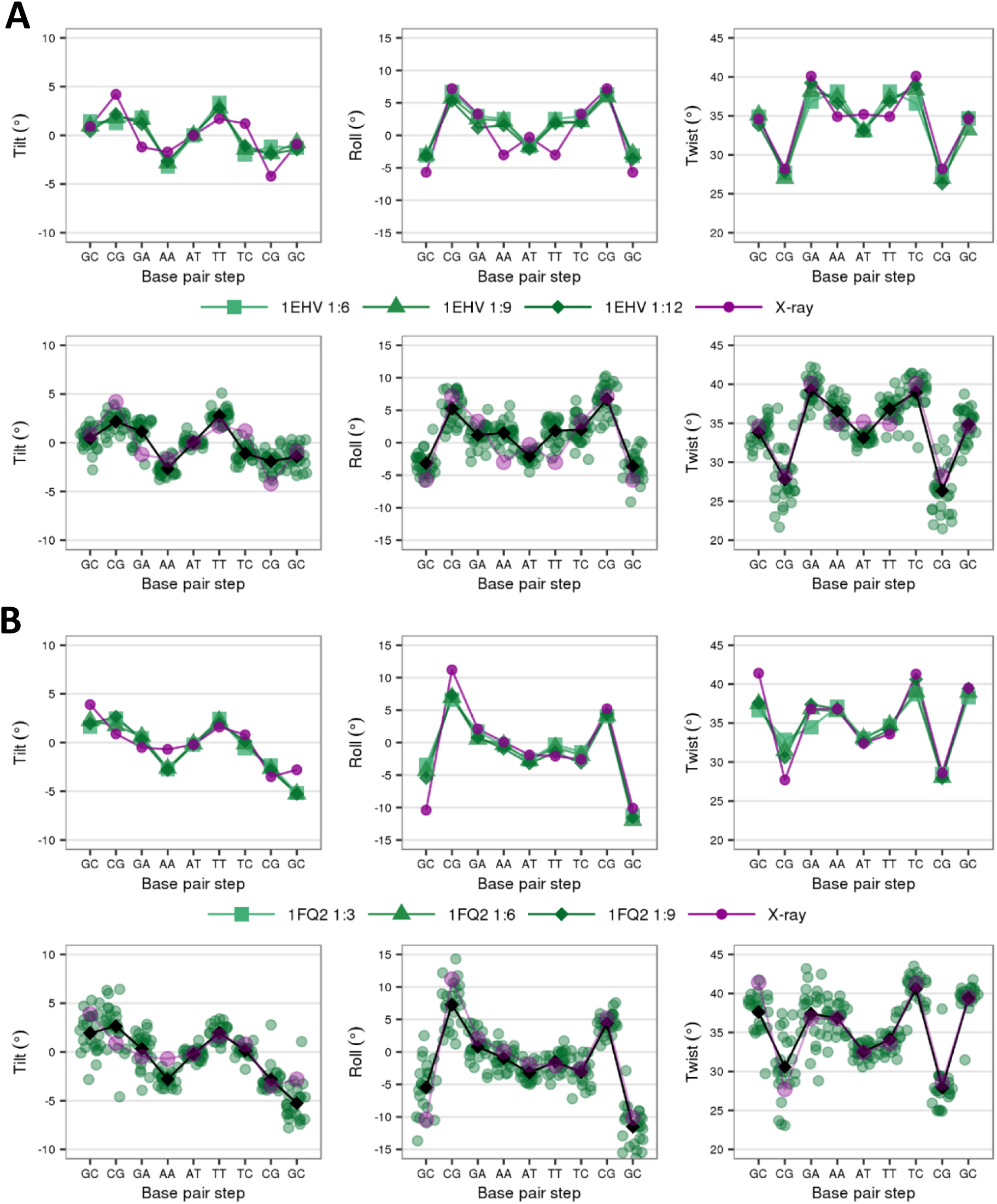
Rotational helical parameters (roll, twist, tilt) (given for all base pair steps in DDD after removing the ends) for different spermine concentrations in A) 1EHV and B) 1FQ2 systems. The upper panels show average values across all duplexes for varying spermine concentrations (colored green), with the purple line giving values calculated for the X-ray structure. The lower panels show the average values for each duplex in the systems with the highest spermine concentration (shown as green circles), while the average over all duplexes is shown as a black line and the X-ray values in purple.

To follow each ion along the trajectories of each duplex, we used the curvilinear helicoidal coordinate (CHC)^39^ method as implemented in Canion/Curves+, which allowed us to calculate ion populations and concentrations in the inner and outer areas of major and minor grooves of DNA duplexes (see Supp. Methods).^40,41^ We found that the sequence-dependent binding sites observed for K^+^ and the amino groups of spermine were the same in solution and in crystals, although the sequence dependence was weaker in crystals and the cation distributions seem to be fuzzier (Supp. Figure S12).^11^ The comparison of aqueous simulations with and without spermine (left column in Supp. Figure S12) showed that spermine mainly localizes outside the grooves, which was also previously observed in crystal simulations.^19,20,38,42^ Interestingly, the interiors of the grooves are depleted of cations in all crystal simulations compared to the solution phase.

We also analyzed the cation environment in all the individual duplexes present in each crystal. First, a clear similarity was found in the cation distribution in all the duplexes of a given crystal, confirming the robustness of the average results discussed here (see Suppl. Figures S13-15). Second, spermine distribution was similar around the duplexes in simulations of two different space groups (1EHV and 1FQ2), indicating that the ion distribution is independent of the crystal packing (see Supp. Figures S14 and S15). Moreover, spermine mostly preferred the exterior of the central part of the major groove, however, upon entering the groove, spermine was located at non-central sites, which we also observed for solution simulations (see Suppl. Figure S12). Spermine could concentrate up to 12 M around the phosphate groups in the central portion of the duplexes where no DNA intermolecular contacts are present. In contrast, these contacts are particularly pronounced in terminal bases (Supp. Figure S16). In summary, spermine has a tendency to move around the exterior of the duplex without any perceptible sequence dependence (as observed in NMR^43^ and RAMAN^44^ experiments), most likely screening intermolecular phosphate-phosphate repulsion and facilitating crystal packing. Combined with its rare long-term presence within the insides of the grooves, it is not surprising that spermine electron density is seldom captured.

While the temperature for both experiments and MD simulations was in the 284-293 K range for 1EHV, 1FQ2, and 463D (see Supp. Methods), the intermolecular DNA contacts in the crystal froze the duplex structure. DNA’s internal effective temperature was lowered by 200 K (with respect to the thermostat temperature) at the end of the duplexes, which are mostly rigidified, while we observed only ∼50 K decrease in the central portion of the duplex (Supp. Figure S17). Water molecules and SPM are also affected by the crowded and highly negatively charged environment produced by the packing of the backbone phosphate groups of neighboring duplexes. The rescaled diffusion coefficient^34^ of water molecules (Supp. Figure S18 and Supp. Methods), was reduced by one order of magnitude - from 1.70·10^-5^ cm^2^ s^-1^ in solution DNA simulations to 1.67·10^-6^ cm^2^ s^-1^ in crystal simulations. The reduction in water mobility is visible from a 30-K decrease in its effective temperature. Furthermore, water mobility in crystals is anisotropic, with the preference for the z-axis in 463D and 1EHV, and the y-axis in 1FQ2 following the water channels that are created in each crystal due to the orientation of the duplexes (Supp. Figure S19). The effect was even more pronounced for SPM when comparing the 1FQ2 1:6 solution simulations with the 1FQ2 crystal with the same SPM concentration: the diffusion coefficient in the crystal was reduced by three orders of magnitude compared to the solution: from 1.90·10^-6^ cm^2^ s^-1^ to 2.62·10^-9^ cm^2^ s^-1^. Although the mobility of SPM is greatly reduced in the crystal, the lack of specificity in its binding to DNA (extensive and non-specific binding outside the major groove) makes its detection in the experimental electron densities very difficult (see Suppl. Figure S13). As expected, the diffusion of water and SPM in the crystal was correlated and exhibited an SPM-concentration dependent effect as shown in Supp. Figure S20.

### How do the structural and dynamical properties of DNA crystals compare with solution?

We computed the global RMSD of MD simulations (crystal and solution) with respect to the experimental crystal configuration (Table 2), confirming that the average MD structure in solution deviates much more from the crystal configuration (with larger deviations in the backbone) than the average structure from the crystal MD simulations,^17^ confirming that MD simulations are able to capture the effect of the crystal environment on the DNA. Irrespectively of the original conformation present in the crystal, when we transferred the duplexes to solution, all the simulations converged to the same ensemble (Supp. Figure S21 and S22), showing a strong consensus and the lack of memory in the simulations. Moreover, as confirmed by WAXS experiments,^38^ SPM has a negligible effect on the structure of DNA in aqueous solution (Supp. Figure S21 and S22). As shown in Figure 5, the solution simulations were generally able to sample the X-ray values within the first standard deviation, indicating that the overall structures in solution and in the crystal are very similar. However, a close inspection shows small local discrepancies between the average MD solution simulations and the X-ray values at several bps, although these differences (Supp. Figure S23 and S24) tend to disappear when average properties of the entire duplex are computed.^11^

**Table 2.**
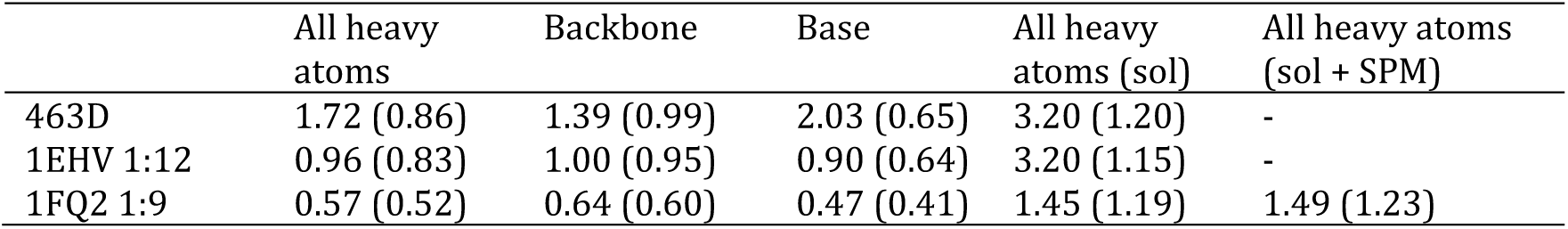
Root mean square deviations (RMSD, Å) from the deposited crystal structures for 463D, 1EHV, and 1FQ2. The statistics in each box are heavy atoms, backbone atoms, and base atoms RMSD of average MD structure with respect to the crystal structure. The values in parentheses exclude the terminal residues.

|  | All heavy atoms | Backbone | Base | All heavy atoms (sol) | All heavy atoms (sol + SPM) |
| --- | --- | --- | --- | --- | --- |
| 463D | 1.72 (0.86) | 1.39 (0.99) | 2.03 (0.65) | 3.20 (1.20) | - |
| 1EHV 1:12 | 0.96 (0.83) | 1.00 (0.95) | 0.90 (0.64) | 3.20 (1.15) | - |
| 1FQ2 1:9 | 0.57 (0.52) | 0.64 (0.60) | 0.47 (0.41) | 1.45 (1.19) | 1.49 (1.23) |

**Figure 5.**
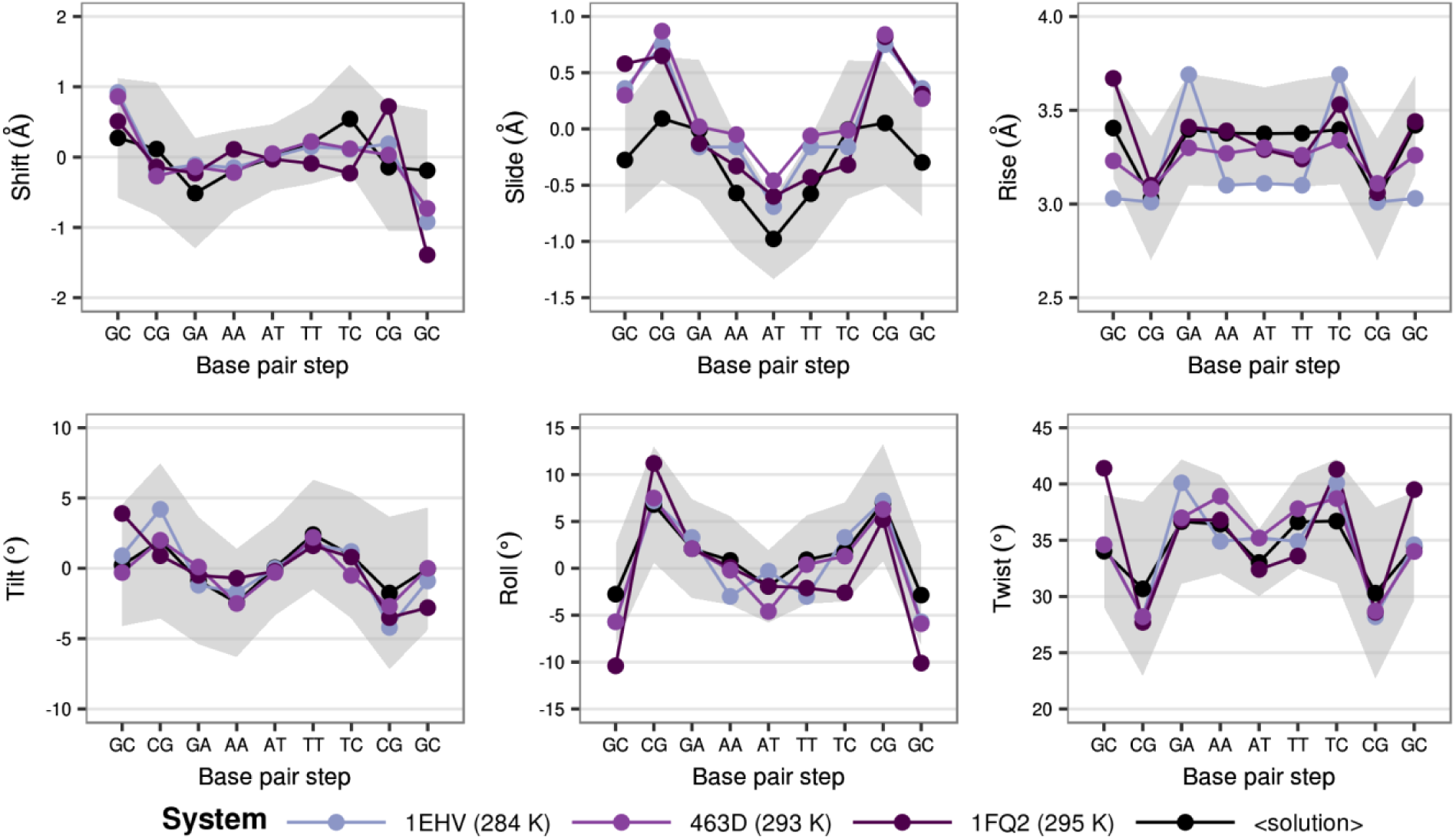
Translational (shift, slide, rise) and rotational (roll, twist, tilt) helical parameters (given for all base pair steps in DDD after removing the ends) for the solution simulations of single duplexes starting from different X-ray structures. The average values across the simulations are shown in black, while the average standard deviations are shown as grey ribbons. Values calculated from X-ray models are shown as colored lines.

We also analyzed how crystal modifies the flexibility and dynamical behavior of the duplex. As expected, DNA in crystals was much more rigid than in solutions, which can be seen from the force constants associated with helical deformations. As expected from helical parameters, the differences are especially pronounced for the terminal residues which are much more mobile in water (Supp. Figure S25). Interestingly, same sequence-dependent pattern of flexibility is found for the crystal and solution simulations -for all degrees of freedom and for all studied crystals. This suggests that the strong reduction of terminal movements is a consequence of the specific DNA packing required for crystallization. The analysis of the essential dynamics (see Methods) of DNA in solution and in the crystal indicates a high degree of similarity: 93-98% for solution *vs* solution simulations, while the similarity decreases to 78-95% for crystal *vs* crystal MD comparisons, and to 75-84% when comparing crystal to solution (Supp. Figure S26). This finding suggests that the essential dynamics of the duplexes in the crystals slightly differs from the one in solution, and that the variability in the type of movements between different crystal lattices is larger than in solution. These results indicate that the small structural discrepancies in the average helical parameters, discussed above, arose from dynamic, rather than static, properties of the DNA. However, they also suggest a reasonable maintenance of the intrinsic deformation pattern of DNA in crystals, which would explain how processes which require deformation of the DNA can be reproduced (on longer timescales) in crystal environments, e.g., binding of small molecules in soaking experiments.^45,46^

We finally focused our analysis on two paradigmatic structural polymorphisms that seem to be causally connected in solution, namely the backbone BI/BII polymorphism (related to ε and ζ torsions), and the bimodal behavior of the twist helical parameter observed for d(CpG) steps.^11,40,47–49^ BII populations are detected in both crystal and solution simulations, but the base pair steps for which such minor populations were observed are different (Supp. Figure 27). In agreement with the experiments (X-ray and 1H/31P-NMR),^50^ BII state was observed less frequently for d(CpG) and d(GpA) steps in the crystal than in the solution simulations, while the opposite was observed for d(GpC) steps. Quite similar results were observed for d(ApA) and d(ApT) steps. This would suggest that the differences in BII propensities between X-ray and NMR structures are not spurious effects related to the uncertainties of the refinement process, but a consequence of crystal packing.

Polymorphisms at the base level (i.e. the existence of two stable conformational substates) were much more difficult to detect experimentally.^51^ On the one hand, NOEs often lack the required resolution to introduce dual restraints in the simulation procedure, and on the other hand, “substate-washing-effect” produced by the averaging of structures present in the crystal annihilate minor substates in the final refined structures.^51,52^ As we have individual data for each duplex in the crystal, we can analyze minor polymorphism with high accuracy. Thus, we explored the twist polymorphism at the d(CpG) step for the three simulated crystals and compared the results to the solution trajectories. We found that the bimodal behavior was mitigated in the crystals due to both averaging effects and a lower propensity for the bimodal twist (Supp. Figures 28-30).^40,49^ The two and three substates observed in the 2D density plots of roll vs twist and twist vs tilt, respectively, were all found among the duplexes present in the crystal units, but not simultaneously in a unique duplex as in the solution simulations. This correlated perfectly with the reduced amount of cations found inside the minor groove of d(CpG) steps in the crystals simulations,^40^ and with the smaller mobility of crystal DNA that reduces the kinetics of substate inter-conversion. These results suggest that the solvent environment created around the duplexes in the crowded crystal affects the DNA dynamics and limits its conformational space compared to the free DNA in solution. Therefore, taking extra caution is necessary before using X-ray data to evaluate small-scale polymorphisms which can be crucial for DNA recognition by ligands.

## CONCLUSIONS

Even with recent advances in cryoEM, X-ray crystallography remains a dominant technique in structural biology that shows no signs of slowing down. However, one of its main challenges still lies in obtaining diffractable crystals. Conditions that lead to such crystals are a priori unknown and the knowledge gained in the field has been mainly empirical. Typically, several types of solvent buffers are usually tried out but since the electron density of ions and chemical additives is rarely observed, it has been impossible to know which buffer components finally end up in the crystal and at which concentration. For example, spermine has been used to obtain thousands of DNA crystals, yet the molecular basis of its mechanism of action was mostly unknown until now. Furthermore, atomistic MD simulations were not able to provide relevant information on the intermolecular forces that dominate within the crystal structure, due to issues with obtaining stable crystal simulations on relevant timescales. In this work, we presented for the first time stable multi-microsecond-long MD simulations of crystals of the Drew-Dickerson dodecamer in all the symmetry groups it has been experimentally crystallized in. We achieved this by using the latest generation classical force field and by modeling correctly the solvent environment around DNA duplexes which allowed us to understand the importance of chemical additives in the crystallization buffer. Once stable, an impressive global and local structural agreement was obtained between the modeled and experimental DNA crystals. This allowed us to unravel the interactions and physical properties of water molecules, ions and spermine at the atomic level, giving their quantifiable overview. Water molecules and spermine diffused 10 and 1,000 times more slowly in the crowded crystal environment than in solution, respectively. Most of the spermine molecules, which could concentrate up to 12 M around DNA, were bound to phosphate groups outside of the major groove without a specific sequence dependence. Moreover, we showed that previous discrepancies in local structure between solution simulations and X-ray structures, or the ones between solution NMR and X-ray structures in the description of the backbone polymorphism, were mainly due to packing effects that modify the solvent-solute interactions, affecting the dynamics of the DNA and limiting its conformational space compared to its free behavior in solution. Our work expands the limits of the field and opens the door to further systematic studies of the effect of crystallization conditions on the obtained structures. In the near future, MD simulations of crystals could be used to anticipate and predict the effects of a given additive, reducing the number of trial and error iterations usually involved in the experimental obtainment of stable DNA crystals.

## SUPPLEMENTARY DATA

Further details on the setup and analysis of the trajectories, and additional results are provided in the Supplementary Methods, Figures and Tables respectively.

## ACKNOWLEDGEMENTS

M.O. is an ICREA (Institucio Catalana de Recerca i Estudis Avançats) academia researcher. P.D.D. is a PEDECIBA (Programa de Desarrollo de las Ciencias Ba sicas) and SNI (Sistema Nacional de Investigadores, Agencia Nacional de Investigacio n e Innovacio n, Uruguay) researcher.

